## Supplementary material for "Multiple origins of dorsal ecdysial sutures in trilobites and their relatives": All supplementary figures

This file includes:

1. **Supplementary figures (Figs. S1–9)**

### 1. Supplementary figures

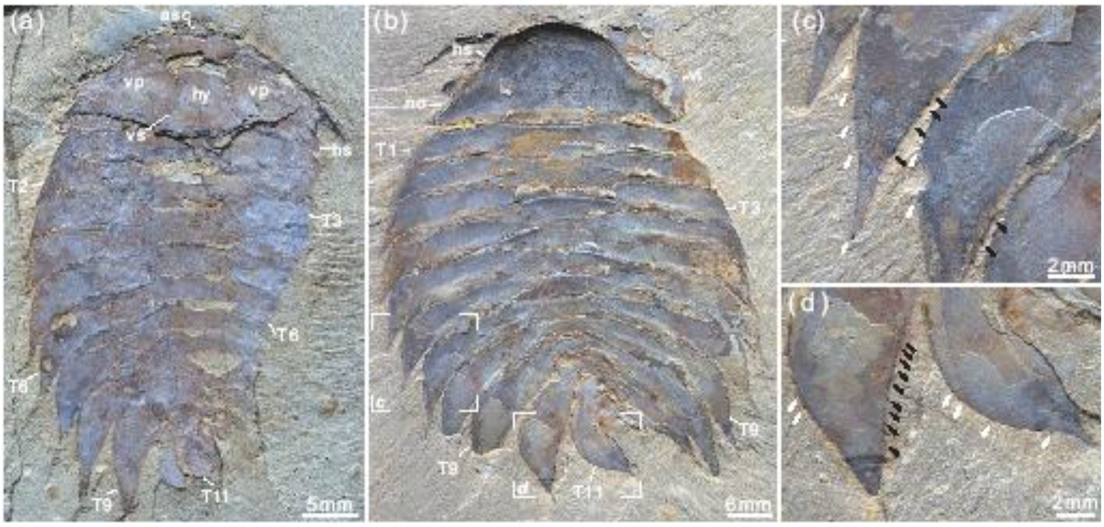


**Fig. S1** ***Acanthomeridion serratum*** **from the Cambrian Stage 3 Chengjiang Biota.** (a) CJHMD 00052b articulated specimen preserved hypostome, librigena-like ventral plates, and 11 tergites with pleural spines. (b) CJHMD 00053b complete specimen with head, ventral plate, and 11 tergites. (c, d) Marginal spines (arrows) on the tergites. Abbreviations: ant, antenna; ca, post-antennal appendage beneath head; dbl, doublure; ds, dorsal suture; en, endopod; ex, exopod; ey, eye; es, eyestalk; gut, digestive tract; hs, head shield; hy, hypostome; lam, lamellae; no, notch; p*n*, podomere *n*; R, right; T*n*, tergite *n*; ts, terminal spine; vp, ventral plate.

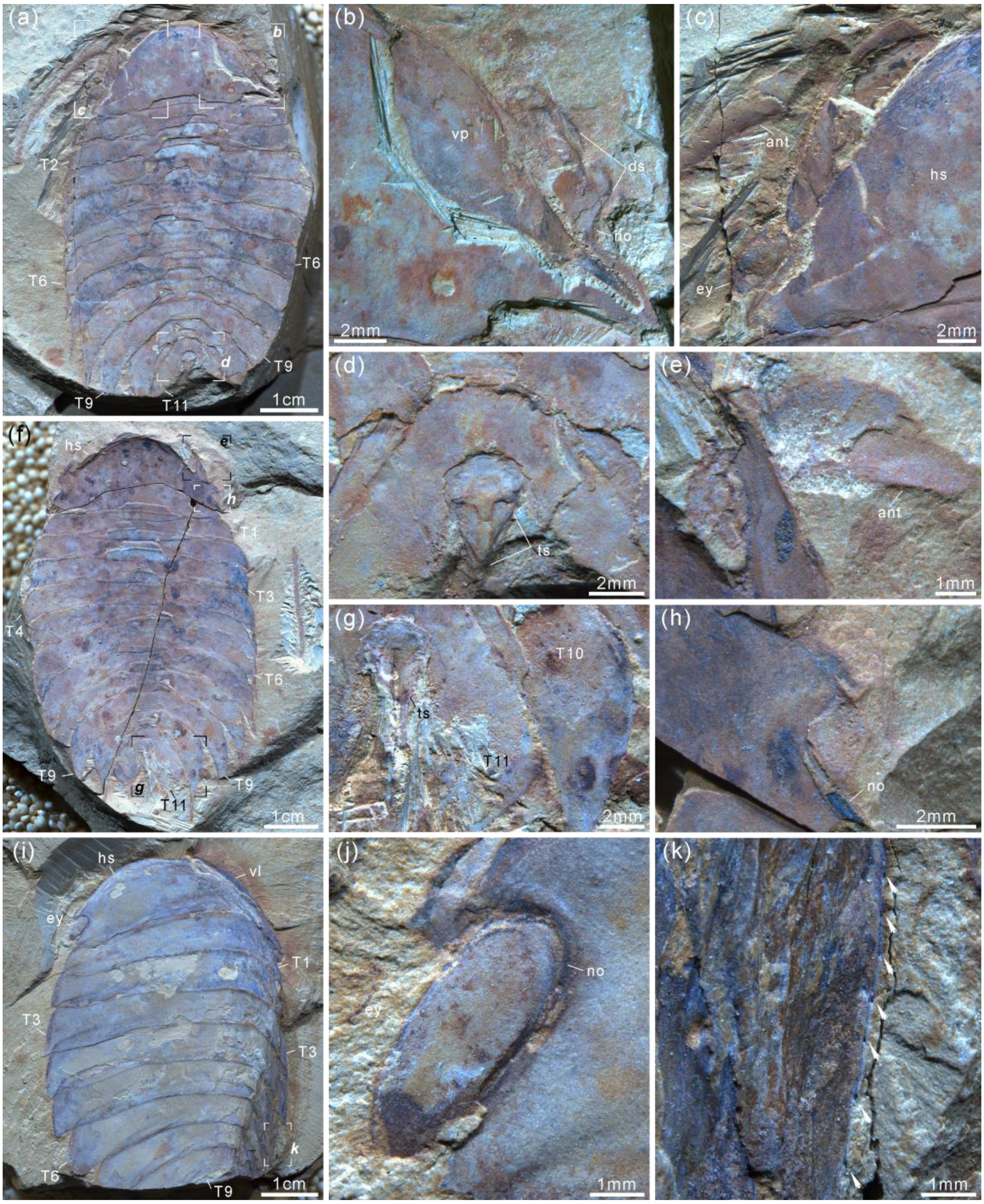


**Fig. S2** ***Acanthomeridion serratum*** **from the Cambrian Stage 3 Chengjiang Biota.** (a) YRCP 0016a, articulated individual with antenna, ventral plates, and 11 tergites. (b) Details of ventral plate and notch. (c) Close-up of antenna and stalk eye. (d, g) Terminal spine and its joint. (e) attachment of antenna. (f) Counterpart of label (a), YRCP 0016b. (h) Detail of notch. (i–k) YRCP 0017, preserved head, compound eye, 9 tergites, and marginal spines on tergite. For abbreviations, see Fig. S1.

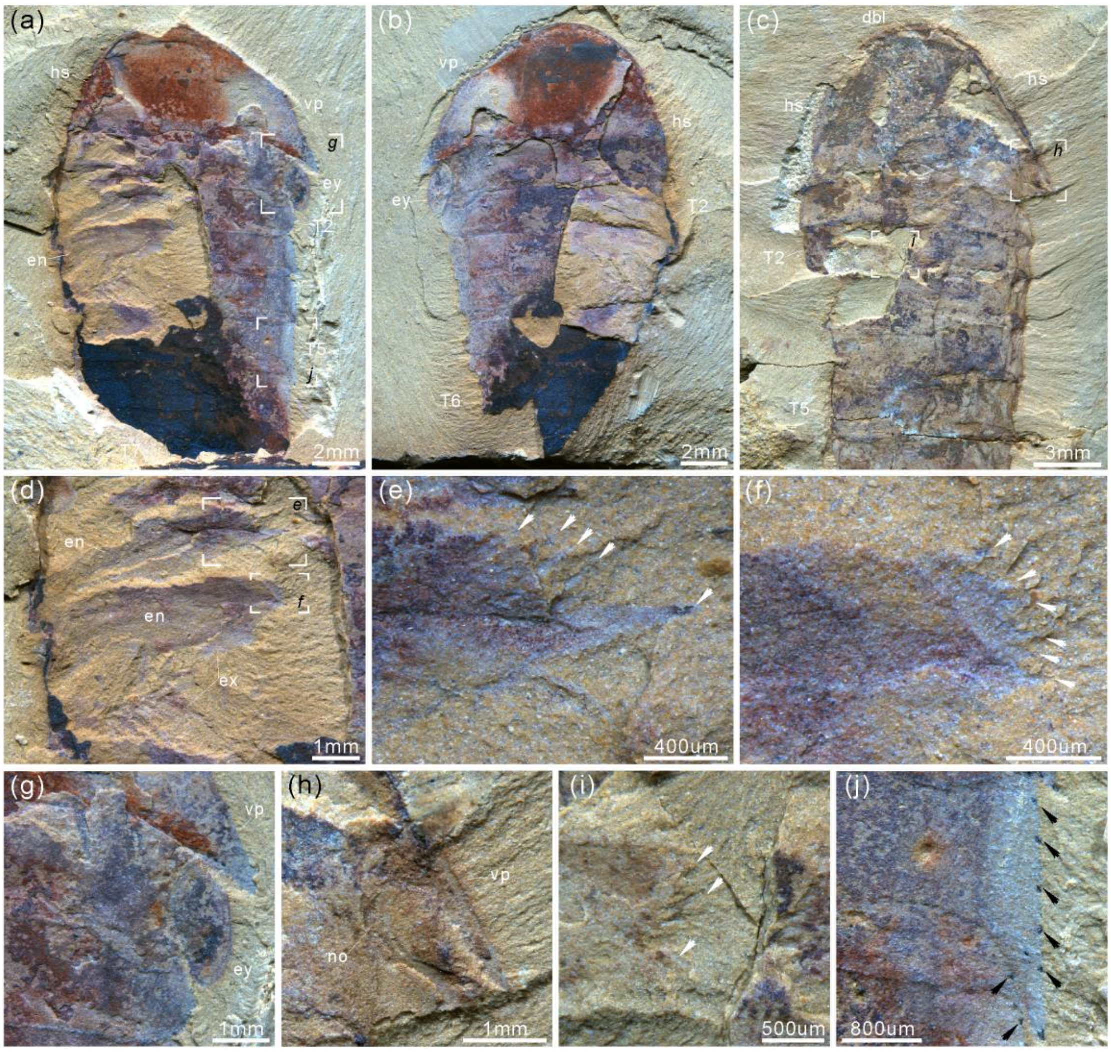


**Fig. S3** ***Acanthomeridion serratum*** **from the Cambrian Stage 3 Chengjiang Biota.** (a) CJHMD 00056a, articulated specimen preserved ventral plates, gnathobases of limbs, endopods, and exopod. (b) CJHMD 00056b. (c) CJHMD 00057, showing the anterior sclerite, head, gnathobases, and 5 tergites. (d) details of gnathobases, endopods, and exopod. (e, f) Close-up of gnathobases. (g, h) Details of elliptical eye and ventral plate. (i) Close-up of gnathobases. (j) Details of marginal spines on tergite. For abbreviations, see Fig. S1.

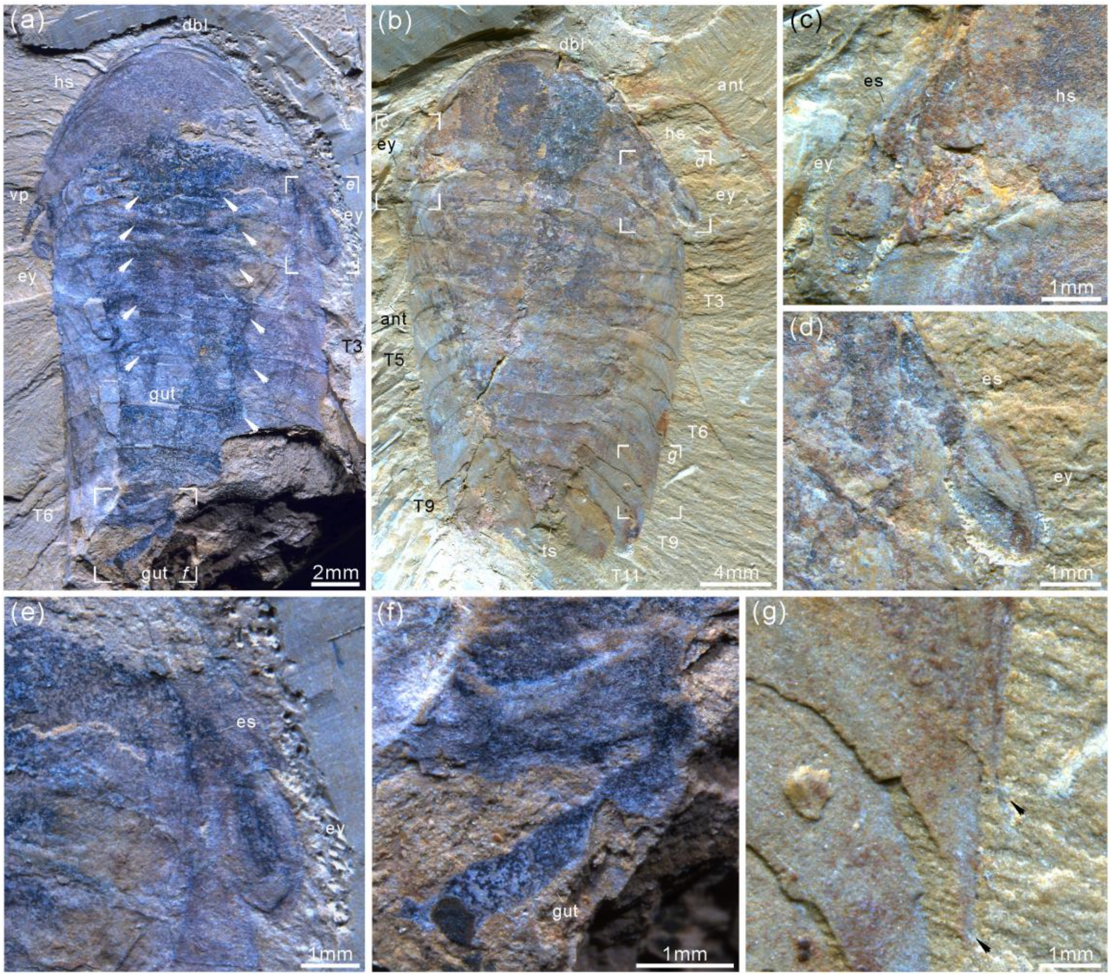


**Fig. S4** ***Acanthomeridion serratum*** **from the Cambrian Stage 3 Chengjiang Biota.** (a) CJHMD 00058, articulated specimen with head, spine of ventral plate, stalked eyes, gut, midgut diverticulae (white arrows), and 6 tergites. (b) YRCP 0018, complete individual with long antennae, compound eyes, and 11 tergites. (c–e) Details of stalked eyes and the eyestalks. (f) Close-up of the gut. (g) Pleural spines of T7 and T8. For abbreviations, see Fig. S1.

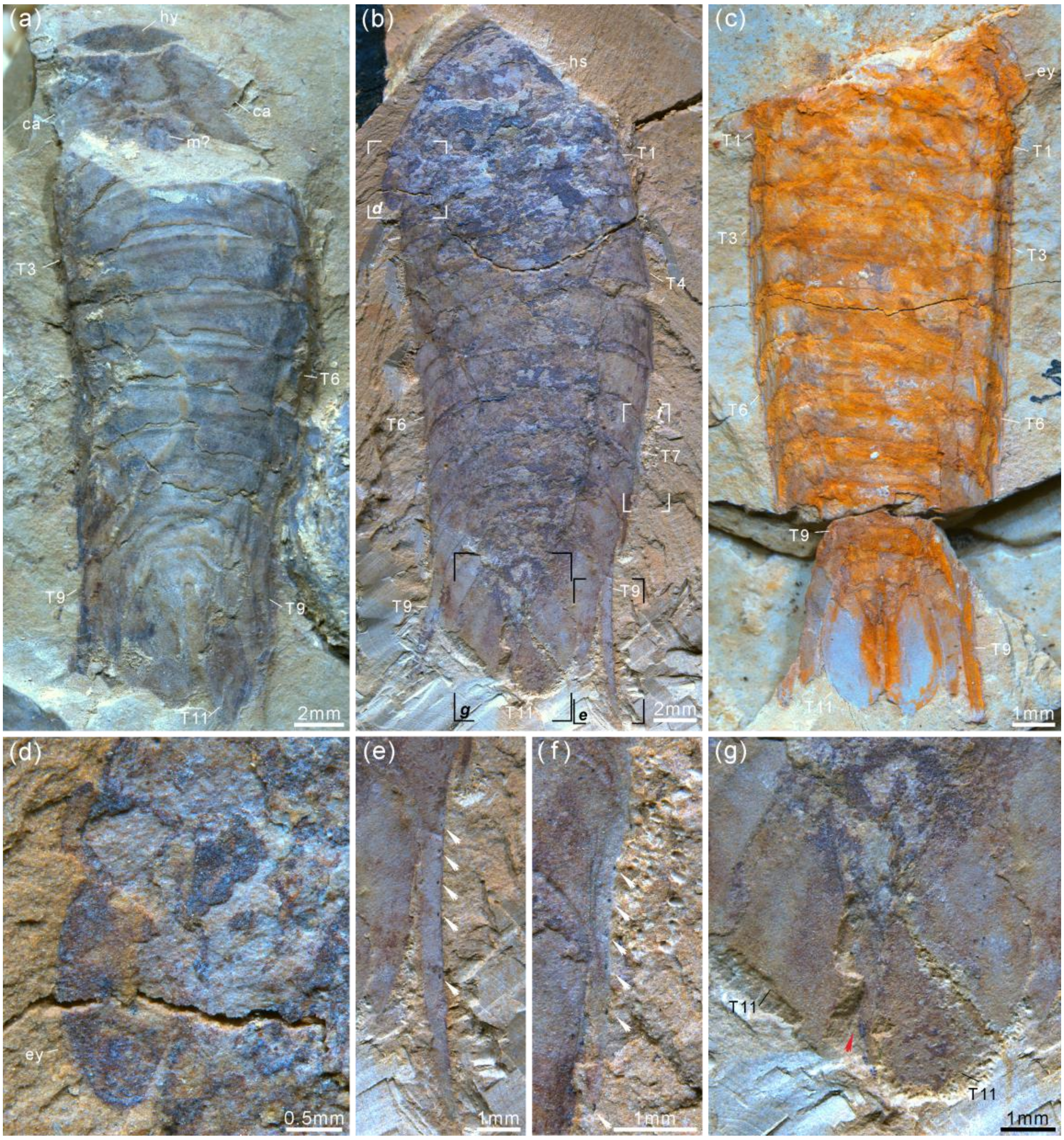


**Fig. S5 *Acanthomeridion serratum* from the Cambrian Stage 3 Chengjiang Biota.** (a) CJHMD 00059, articulated individual with hypostome, cephalic appendages, possible mouth, and 11 tergites. (b) YRCP 0019, complete individual with stalked eyes, long spines on T9, and 11 tergites. (c) YRCP 0020, articulated specimen with compound eyes and 11 tergites. (d) Elliptical eye. (e) Marginal spines (white arrows) on the long pleural spine of T9. (f) Marginal spines (white arrows) on the pleural spine of T7. (g) Paddle-like structure (red arrow). For abbreviations, see Fig. S1.

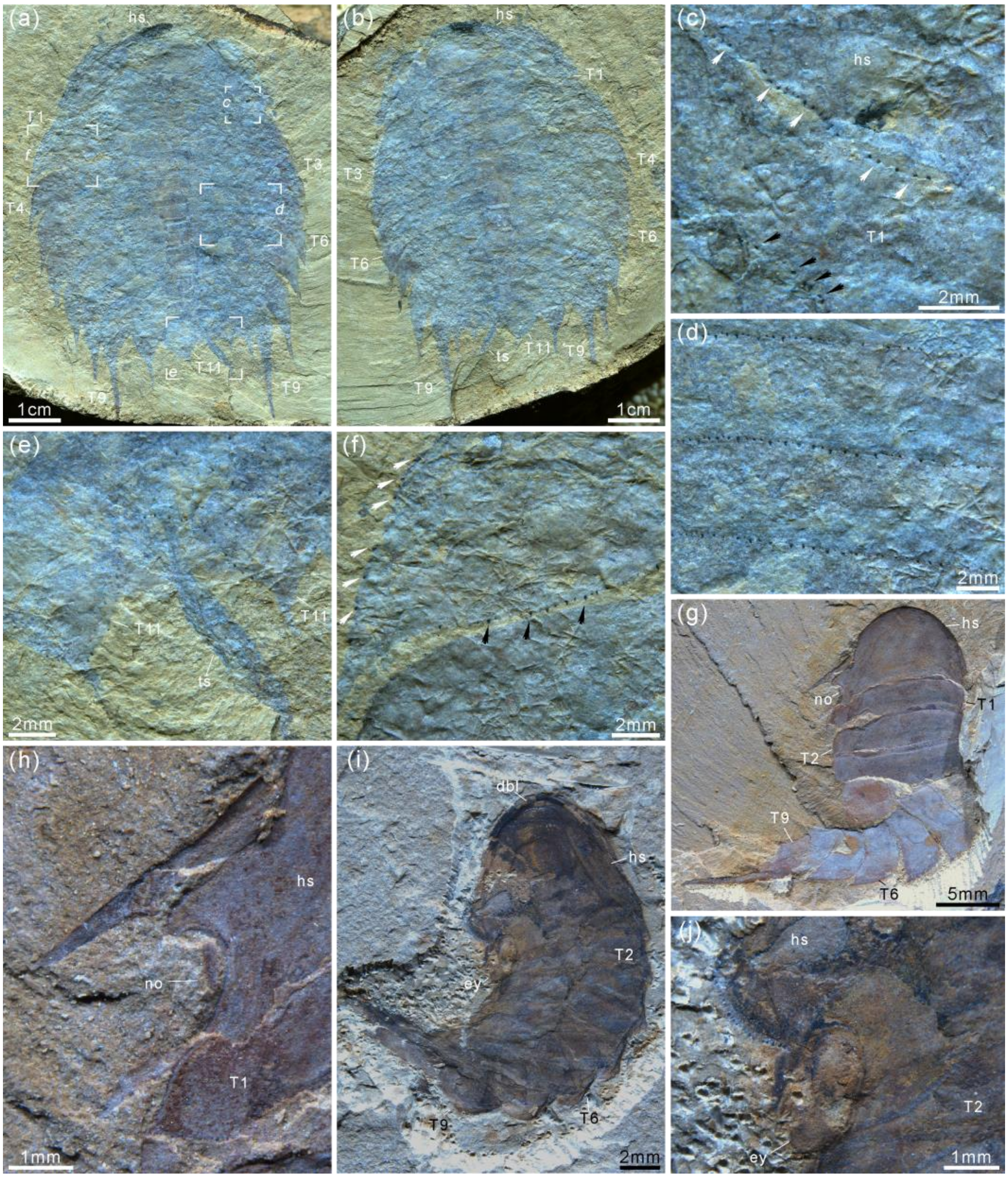


**Fig. S6** ***Acanthomeridion serratum*** **from the Cambrian Stage 3 Chengjiang Biota.** (a) CJHMD 00060a, complete individual with head, 11 tergites, well-developed pleural spines, and terminal spine. (b) Counterpart of CJHMD 00060b. (c) Marginal spines on the posterior margin of head (white arrows) and T1 (black arrows). (d) Marginal spines on the posterior margin of tergites. (e) Pleural spines and marginal spines of T11, terminal spine. (f) Marginal spines on the posterior and lateral margins of tergite. (g, h) CJHMD 00061, articulated juvenile individual with head, notch, and 9 tergites. (i, j) CJHMD 00062, juvenile individual with stalked eyes. For abbreviations, see Fig. S1.

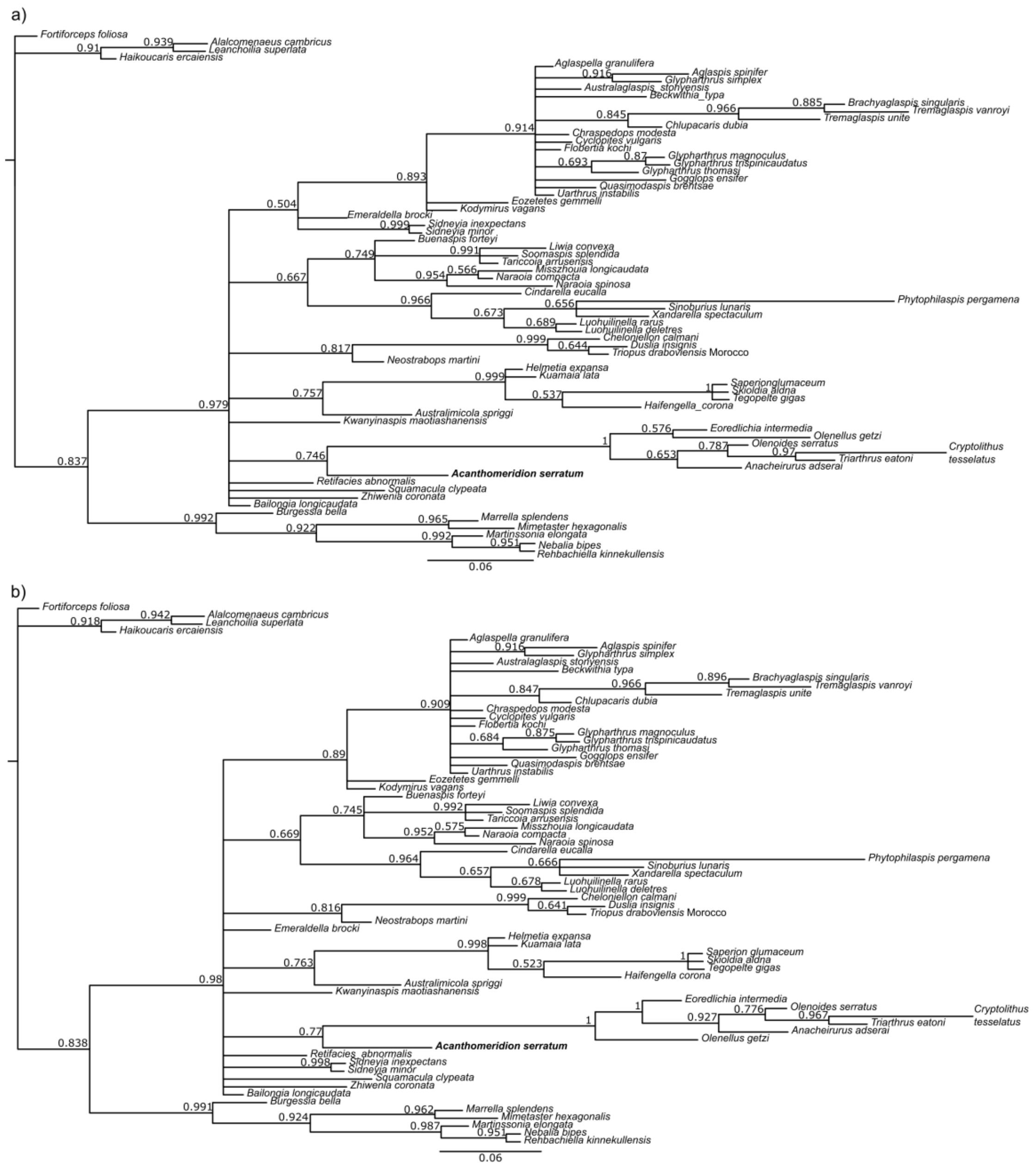


**Fig. S7 Results of phylogenetic analyses using Bayesian inference, with the ventral plates of *Acanthomeridion* treated as homologous to the librigenae of trilobites.** (a) unconstrained analysis. (b) clade comprising *Anacheirurus adserai, Cryptolithus tesselatus, Eoredlichia intermedia, Olenoids serratus, Triarthrus eatoni* constrained. Values at nodes are posterior probabilities.

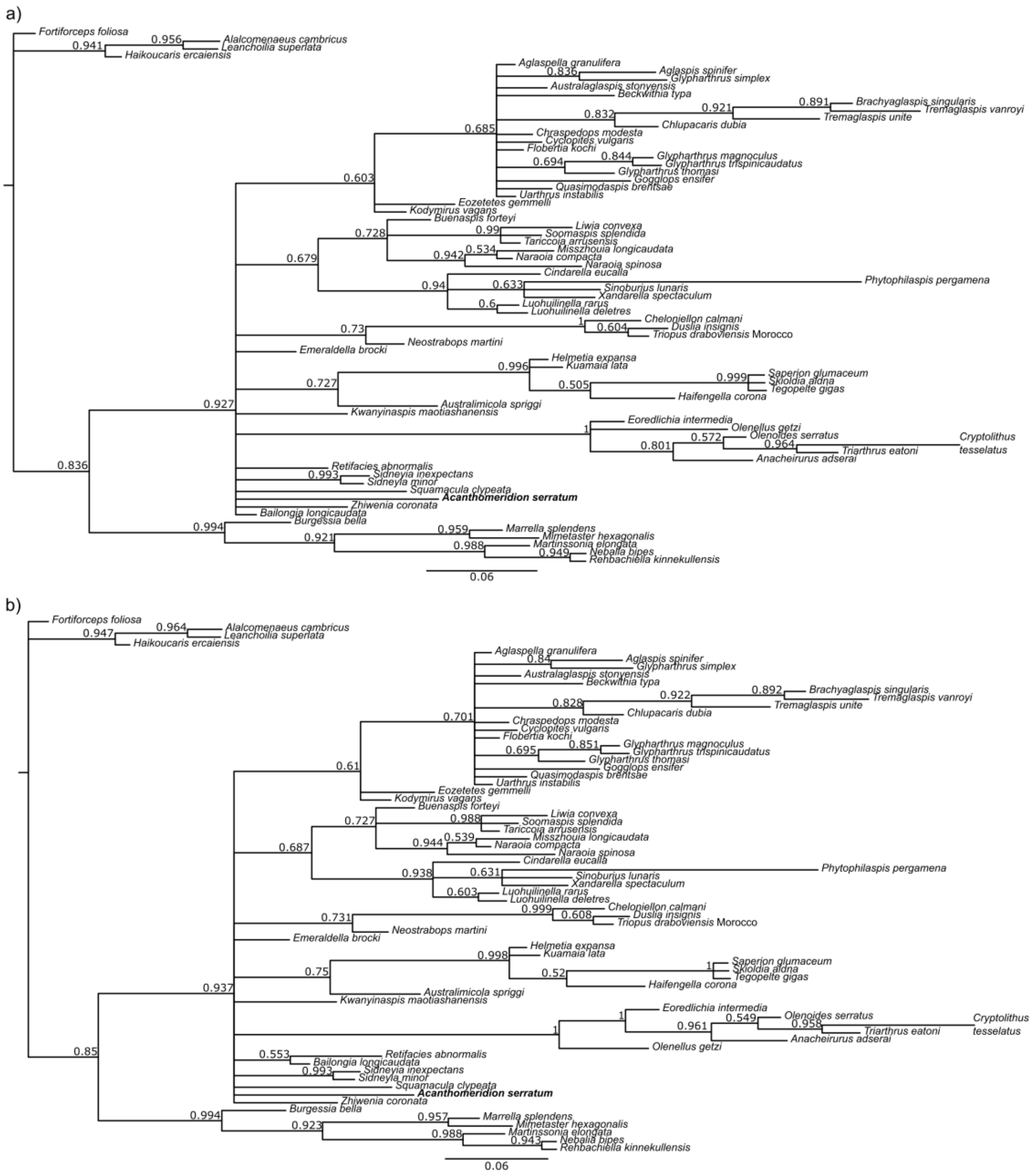


**Fig. S8 Results of phylogenetic analyses using Bayesian inference, with the ventral plates of *Acanthomeridion* treated as homologous to cephalic doublure of trilobites.** (a) unconstrained analysis. (b) clade comprising *Anacheirurus adserai, Cryptolithus tesselatus, Eoredlichia intermedia, Olenoids serratus, Triarthrus eatoni* constrained. Values at nodes are posterior probabilities.

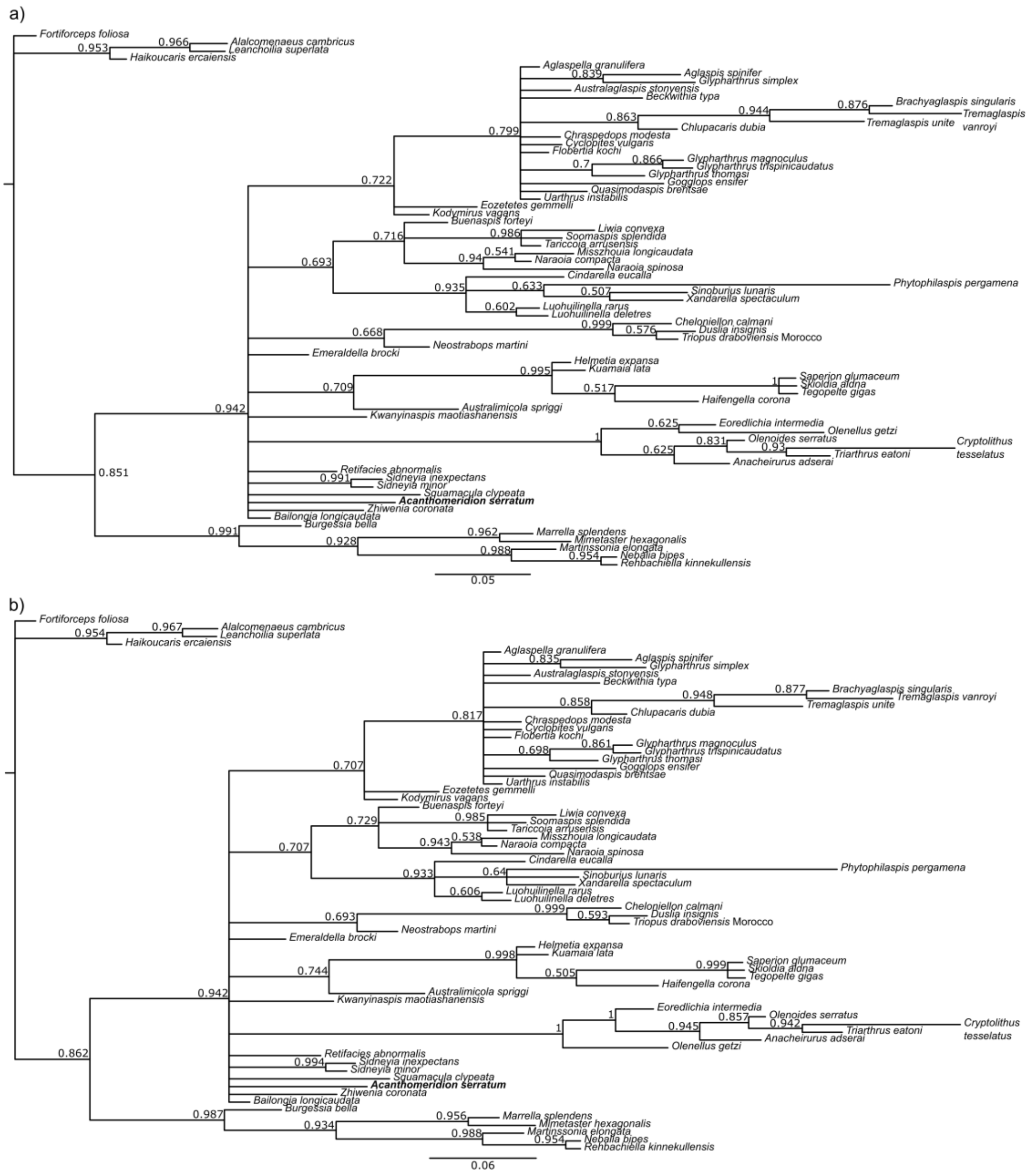


**Fig. S9 Results of phylogenetic analyses using Bayesian inference, with the ventral plates of *Acanthomeridion* treated as non homologous to any cephalic feature of trilobites.** (a) unconstrained analysis. (b) clade comprising *Anacheirurus adserai, Cryptolithus tesselatus, Eoredlichia intermedia, Olenoids serratus, Triarthrus eatoni* constrained. Values at nodes are posterior probabilities.
